# CRSIPR-A-I: a webtool for the efficacy prediction of CRISPR activation and interference

**DOI:** 10.1101/2021.12.02.470943

**Authors:** Xiao Zheng, Jiajun Cui, Yixuan Wang, Jing Zhang, Chaochen Wang

## Abstract

CRISPR-based gene activation (CRISPRa) or interference (CRISPRi) are powerful and easy-to-use approaches to modify the transcription of endogenous genes in eukaryotes. Successful CRISPRa/i requires sgRNA binding and alteration of local chromatin structure, hence largely depends on the original epigenetic status of the target. Consequently, the efficacy of the CRISPRa/i varies in a wide range when applied to target different gene loci, while a reliable prediction tool is unavailable. To address this problem, we integrated published single cell RNA-Seq data involved CRISPRa/i and epigenomic profiles from K562 cells, identified the significant epigenetic features contributing to CRISPRa/i efficacy by ranking the weight of each feature. We further established a mathematic model and built a user-friendly webtool to predict the CRISPRa/i efficacy of customer-designed sgRNA in different cells. Moreover, we experimentally validated our model by employing CROP-Seq assays. Our work provides both the epigenetic insights into CRISPRa/i and an effective tool for the users.

## 1 Introduction

The precise control of gene expression is central to cell fate determination in every living organism. To artificially alter the cell state, genetic and epigenetic tools have been developed to manipulate gene expression. The CRISPR/Cas9 technology is an efficient tool for genome editing based on the duplex of DNA-binding proteins (Cas9 endonucleases) and the single guide RNA (sgRNA) (Kampmann, 2018). With this technology, a genome locus could be accurately targeted, cleaved and mutated through non-homologous end joining (NHEJ) or homologous recombination (HR).

Beyond genetic editing, an advanced application of CRISPR has been developed to alter the expression of target genes instead. The CRISPR activation/interference (CRISPRa/i) technology, uses a nuclease-deficient form of Cas9 (dCas9) fused with a transcriptional effector to activate or repress the expression of the gene locus navigated by sgRNA. With CRISPRa/i, one can modulate the expression of specific genes without changing the DNA sequence. dCas9-VP64 and dCas9-P300 have been widely utilized in CRISPRa. VP64, a transcription activator, facilitates the deposition of activated histone marks and chromatin accessibility to activate gene transcription (Beerli et al., 2000). P300 catalyzes histone H3 lysine 27 acetylation (H3K27ac) causing robust transcriptional activation at diverse regions of target genes, including promoter, proximal enhancers and distal enhancers (Hilton et al., 2015). When dCas9 fused to KRAB repression domain, gene transcription is repressed through steric hindrance or nuclease-based disruption (Kearns et al., 2015). However, all CRISPR-Cas9-based applications still face the challenge of designing optimal sgRNA to improve targeting and the consequent genome editing or regulation.

Currently, at least 30 web-based sgRNA design tools are in use, which are primarily sequence-based, maximizing on-target activity while minimizing off-target activity (Hanna and Doench, 2020). These tools provide a comprehensive sequence analyses, such as the GC content, repetitive sequences, secondary structure of sgRNA, sites for restriction endonucleases, nucleotide identity and thermodynamics. Off-target score is commonly calculated by mismatch count, and Cutting Frequency Determination (CFD) score, etc. On-target scoring algorithms include Rule Set 2, Rule Set 1, and Moreno-Mateos etc. (Hanna and Doench, 2020). However, in addition to the sequence, the genomic position of the sgRNA is also important for the success of CRISPRa and CRISPRi. It was reported that the efficacy of CRISPRi reduces with increasing distance of sgRNA to the transcription start site (TSS) (Radzisheuskaya et al., 2016; Horlbeck et al., 2016).

In eukaryotes, the endogenous transcription occurs with the coordination of the transcriptional factors and permissive epigenetic modulators, which opens the chromatin to facilitate general transcription machinery binding to the DNA; while gene silencing is mediated by co-repressors and restrictive epigenetic modulators, which retain the chromatin closed (Li et al., 2007; Kobayashi and Kurumizaka, 2019). As a result, the difficulty of artificial activation/repression of a silent/active gene by CRISPRa/i is largely determined by how closed/open the chromatin was, that is characterized by epigenetic marks. The active genes are marked by permissive epigenetic marks, including acetylation of histones, histone H3K4 methylation, H3K36 methylation, etc. On the other hand, restrictive epigenetic marks, like DNA methylation, H3K27me3 and H3K9me2/3 are enriched on silent genes (Kouzarides, 2007; Morgan and Shilatifard, 2020). Therefore, we reasoned that the epigenetic features of the target genes could serve as CRISPRa/i predictors. In this study, we took advantage of high-throughput CRISPRa/i coupled with single cell RNA-Seq and ensemble machine learning models, and established a tool to predict the efficacy of CRISPRa/i using epigenetic marks.

## 2 Method

### 2.1 Data preparation

#### 2.1.1 Epigenetic data retrieval and processing

Human (K562, H1, H9 and IMR-90 cell line) and mouse (MEL and ES cell line) reference epigenomes data were downloaded from ENCODE (http://www.encodeproject.org) (Supplementary information). Raw data were sorted and then tag directories were created with the HOMER suite for further peak annotation.

To identify the real cleavage site of each sgRNA in each dataset, sgRNA sequences were aligned to the genome via BLAT software. Window sizes of each epigenetic feature were set based on their different modification scopes, including 100bp (DNase), 200bp (H2AFZ, H3K4me3), 500bp (H3K4me1, H3K4me2, H3K9ac, H4K20me1, H3K27ac, H3K36me3, H3K79me2) and 1000bp (H3K9me3, H3K27me3), taking the cleavage site as the center. Then, peaks in the window sizes were annotated with the HOMER suite for model training.

#### 2.1.2 Single-cell data retrieval and processing

For model training, the single-cell sequencing matrices after pooled CRISPR screening experiments were downloaded from GSE146194 (GSM4367985 for CRISPRi, and GSM4367986 for CRISPRa experiments) which were generated by (Replogle et al., 2020). We first defined the background as the cell group without any sgRNA induced, and calculated the log2 mean expression of each target gene (with pseudocount = 100). Then, for each sgRNA, the log2 mean expression of its target gene was calculated (with pseudocount = 100), which was then subtracted by the log value of background of the corresponding gene. Therefore, the activity score of each sgRNA can be expressed as follows:

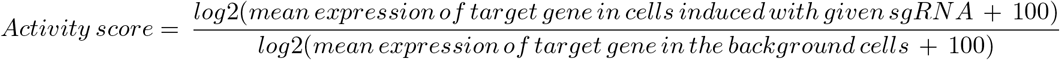

The activity scores for the test data were calculated in the same way. Note that the cells which were expected to contain sgRNAs while the sgRNAs were not detected were excluded before calculating the activity scores.

### 2.2 Wet lab experiment

#### 2.2.1 Cell culture

HEK293T and K562 cells were cultured in Dulbecco’s Modified Eagle’s Medium (DMEM) medium (06-1055-57-1ACS, BI) and RPMI1640 medium (01-100-1A, BI) respectively. The mediums were supplemented with 10% fetal bovine serum (FBS) (FSP500, ExCell Biotech), 1% penicillin/streptomycin (P/S) (03-031-1B, BI). The cell cultures were incubated at 37°C in 5% CO2, and both cell lines were tested for mycoplasma contamination.

#### 2.2.2 SgRNA library preparation

sgRNAs targeting 32 genes were selected from https://www.addgene.org/pooled-library/weissman-human-crispra-v2-subpools/. The sgRNAs were cloned into the CROPseq-Guide-mCherry backbone by Gibson assembly, and amplified in EnduraTM electrocompetent cells (60242-2, Lucigen) at 37°C overnight. Finally, the colonies were harvested, and the plasmids were prepared. The coverage and distribution of the sgRNA library was verified by next-generation sequencing (NGS).

#### 2.2.3 Lentiviruses package and FACS sorting

Lentiviruses was packaged in H3K293T cells with co-transfection of packaging plasmid psPAX2 [Addgene #12260], enveloping plasmid pMD2.G [Addgene #12259] and lentiviral vector (plv-dCas9-VP64-puro, Lenti-CROPseq-Guide-sgRNAs-mCherry) [Addgene #86708]. After 48-72h incubation, the lentiviruses were harvested from the supernatant and added into culture medium. During 2 week’s infection, the cells labeled with Mcaherry were sorted using Fluorescence-Activated CellSorting (FACS) method.

#### 2.2.4 Single cell library construction

Cells were loaded into Microfluidic Chip (GEXSCOPE Single Cell RNA-seq Kit, Singleron Biotechnologies) and processed through Singleron MatrixTM automated single-cell sequencing library construction system for droplet generation. Then, the cells underwent drop lysis followed by reverse transcription. Then, cDNA was isolated and amplified by PCR. Amplified cDNAs then underwent purification, fragmentation, end repair, and A-tailing. Adaptors and sample indices were then added by PCR. Samples were then selected by Qseq100. The resulting library constructs containing P5 and P7 Illumina sequencing adaptors, cell barcode, unique molecular identifier (UMI), genes insert and an i7 sample index. Finally, the resulting scRNA-seq libraries were sequenced on an Illumina scFTD seq instrument.

### 2.3 Model construction

#### 2.3.1 Machining learning model construction

In the project, thirteen regressors were constructed initially, and seven of them were ultimately chosen for CRISPRa sgRNA activity score prediction and three of them were selected for CRISPRi. When constructing models, Scikit-learn was used, the whole process including training, testing and parameter tuning was performed using python3.7.

#### 2.3.2 Web development

The sgRNA design algorithm was derived and modified from (Joung et al., 2017), and the website is shown as follows: https://github.com/fengzhanglab/Screening_Protocols_manuscript/blob/master/design_library.py. a Javaweb was utilized in web development. The UI was built based on JSP and the backstage was connected to R, shell, and python scripts for different information retrieval. A cloud sever is responsible to store the data, run the project and publish the web through Tomcat.

## 3 Result

### 3.1 Overview of workflow

To systematically explore the efficacy of CRISPRa/i in mammalian cells, we pursed a previous study that used single cell transcriptomics to assess the effect of dCas9-VP64 and dCas9-KRAB-mediated gene activation and repression in batch in K562 cells (Replogle et al., 2020). We re-analyzed the data to obtain the expression foldchange corresponding to each sgRNA. Meanwhile, the epigenetic features, including various histone modifications and chromatin accessibility at each locus were extracted and quantified based on the public ChIP-Seq and DNase-Seq data in ENCODE project. Taken the quantified epigenetic features and the distance-to-TSS of sgRNA as input and the foldchange of sgRNA-targeted gene as output, we then employed 10 models (7 for CRISPRa and 3 for CRISPRi) for machine learning (Figure 1A). Eventually, built upon the trained models, a user-friendly web-tool, named “CRISPR-A-I” (http://crispr-ai.net) was deployed to predict the efficacy of CRISPRa or CRISPRi by user-defined sgRNA(s) in selected cells (Figure 1B).

**Figure 1:**
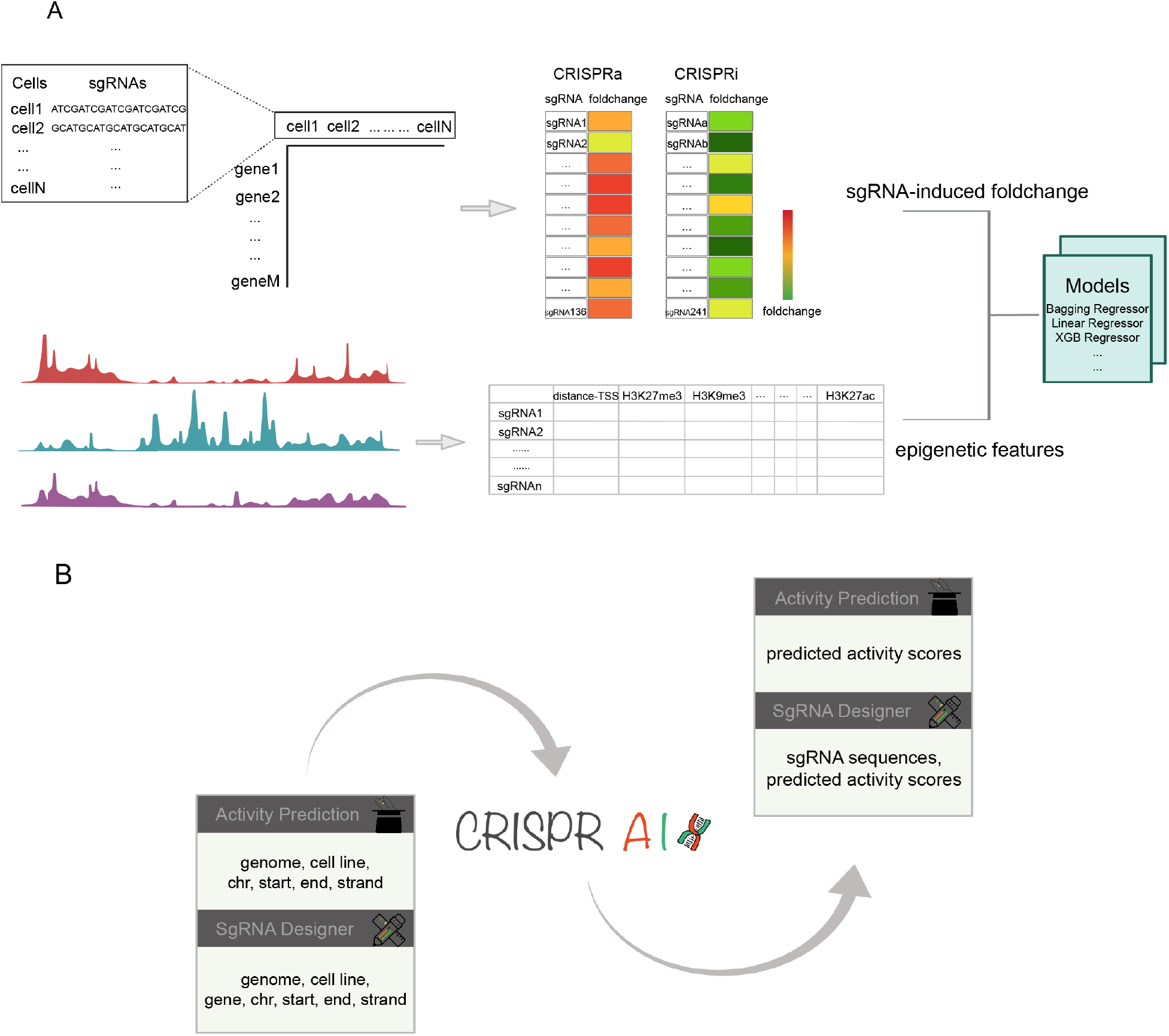
The project overview. (A) Models were trained with epigenetic features as the input and sgRNA-induced foldchanges as the target. Single cell RNA-seq data were analyzed and epigenetic features around each cleavage site were also quantified. (B) Based on the models selected, clients can either predict efficacy of their own sgRNAs or design satisfactory sgRNAs.

### 3.2 The efficacy of CRISPRa/i by different sgRNAs varies in a wide range

377 sgRNAs (136 for CRISPRa and 241 for CRISPRi) targeting 147 genes (55 for CRISPRa and 92 CRISPRi) were included in the train dataset. As expected, the CRISPRa- and CRISPRi-induced gene expression alteration is shifted towards positive or negative in general, respectively. Notably, the log2 foldchanges of the targets ranged from −0.46 to 18.79 in CRISPRa and −5.06 to 12 in CRISPRi, indicating that the efficacy at distinct gene loci varies considerably. Nevertheless, the log2 fold change of genes approximately fitted normal distribution, centralized in the range of (0, 5) in CRISPRa dataset and (−5, 0) in CRISPRi (Figure 2).

**Figure 2:**
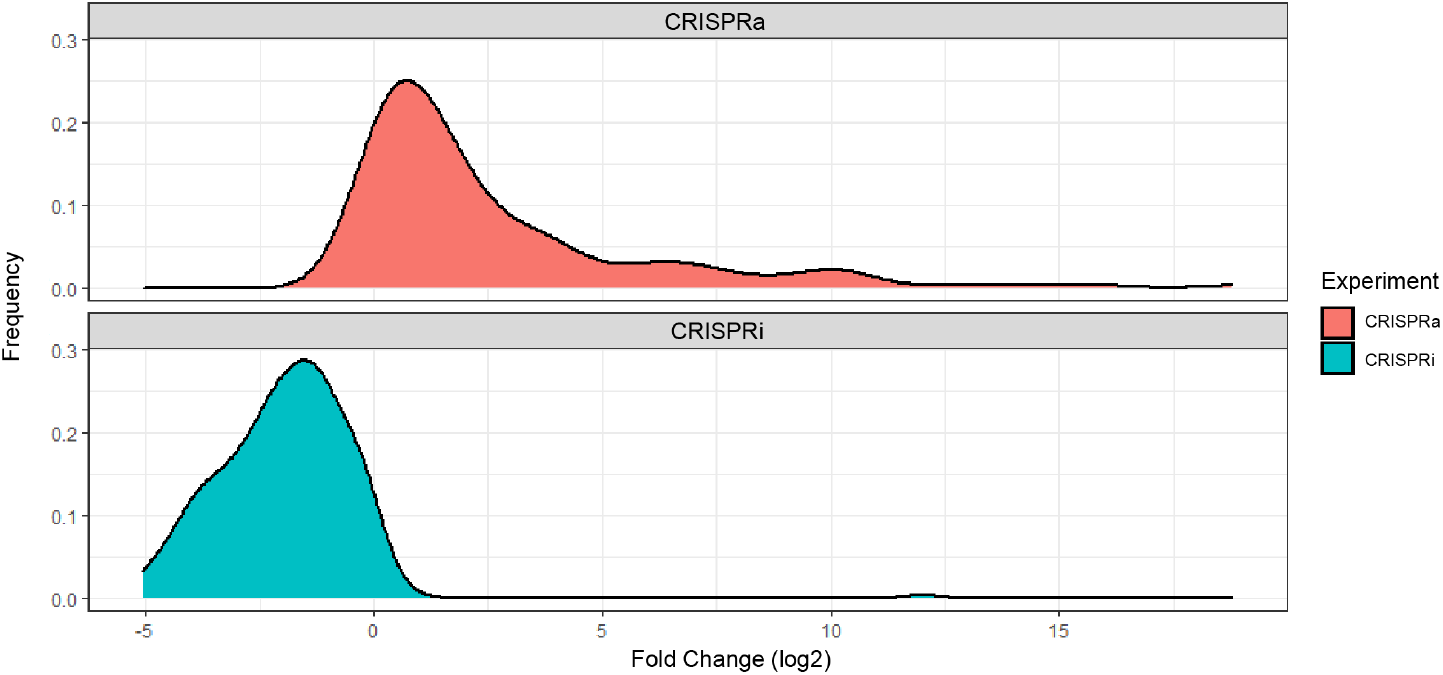
Density plot of log2 fold change of the genes targeted by sgRNAs in CRISPRa and CRISPRi experiments, respectively. The data were generated from CRISPRa/i-based multiplexed single-cell sequencing experiments done by (Replogle et al., 2020)

### 3.3 Mathematical model of CRISPRa/i efficacy by machine learning

Initially, 13 commonly used machine learning models were trained with data consisting of 15 epigenetic features and corresponding activity scores of target genes (log2 fold change). Using the score function from scikit-learn, the coefficient of determination (R2) was calculated to estimate the performance of each model. The results showed that ensemble regressors including bagging and boosting models performed better and had higher R2 scores (R2>0.85) than individual models in CRISPRa (Figure 3A). Eight regression models scoring over 0.9 were selected for CRISPRa.

**Figure 3:**
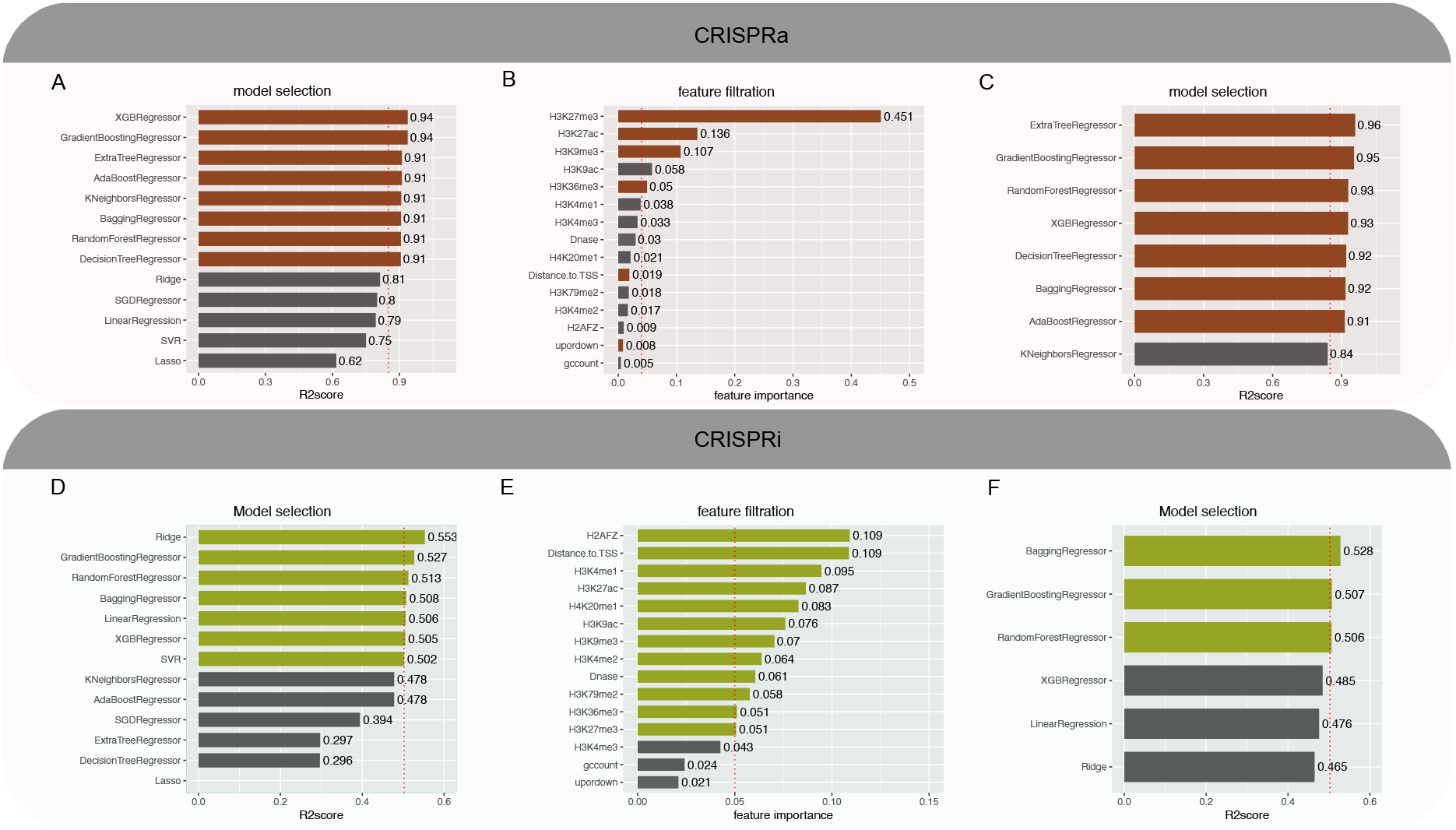
Model & feature selection. (A) Barplot of R2score of indicated models after CRISPRa data training. (B) Barplot of feature importance ranking calculated with sklearn based on the indicated models. (C) Barplot of R2score of indicated models with six sustained features. (D-F) Barplots presenting the R2scores of models and feature importance ranking with CRISPRi data, as in A-C.

Based on selected models, the ranking of feature importance was generated through the ‘feature_importances_’ function in scikit-learn package. Then, the epigenetic features were screened to eliminate redundancy based on both the statistical ordering and biological characteristics (Figure 3B). After feature filtration, six features were left to build the models for CRISPRa prediction again. 7 of 8 models performed better with these features (Figure 3C). Therefore, these models were selected to predict sgRNA activity in CRISPRa eventually. We also applied similar procedure in CRISPRi modeling and 3 models with 12 features were selected lastly (Figure 3D–F).

### 3.4 Cross-validation of prediction models by CROP-Seq

To validate our prediction models, we performed CROP-Seq experiment with dCas9-VP64 mediated CRISPRa containing 132 sgRNAs targeting 33 genes in K562 cells. Post-sequencing analysis shows that most sgRNAs were detected in more than 5 cells (Figure 4A) and more than 80 sgRNAs had over 50 unique aligned reads (Figure 4B), demonstrating a high proportion of cells successfully transduced. By excluding low-quality reads and cells, we collected data from 15780 K562 cells with 60 sgRNAs targeting 25 genes. Clustering analysis of the single-cell transcriptomics showed that cells carrying the same sgRNAs present similar transcription programs, while cells with distinct sgRNAs are clustered separately, elucidating the reproducibility and effectiveness of CRISPRa on their target genes (Figure 4C).

**Figure 4:**
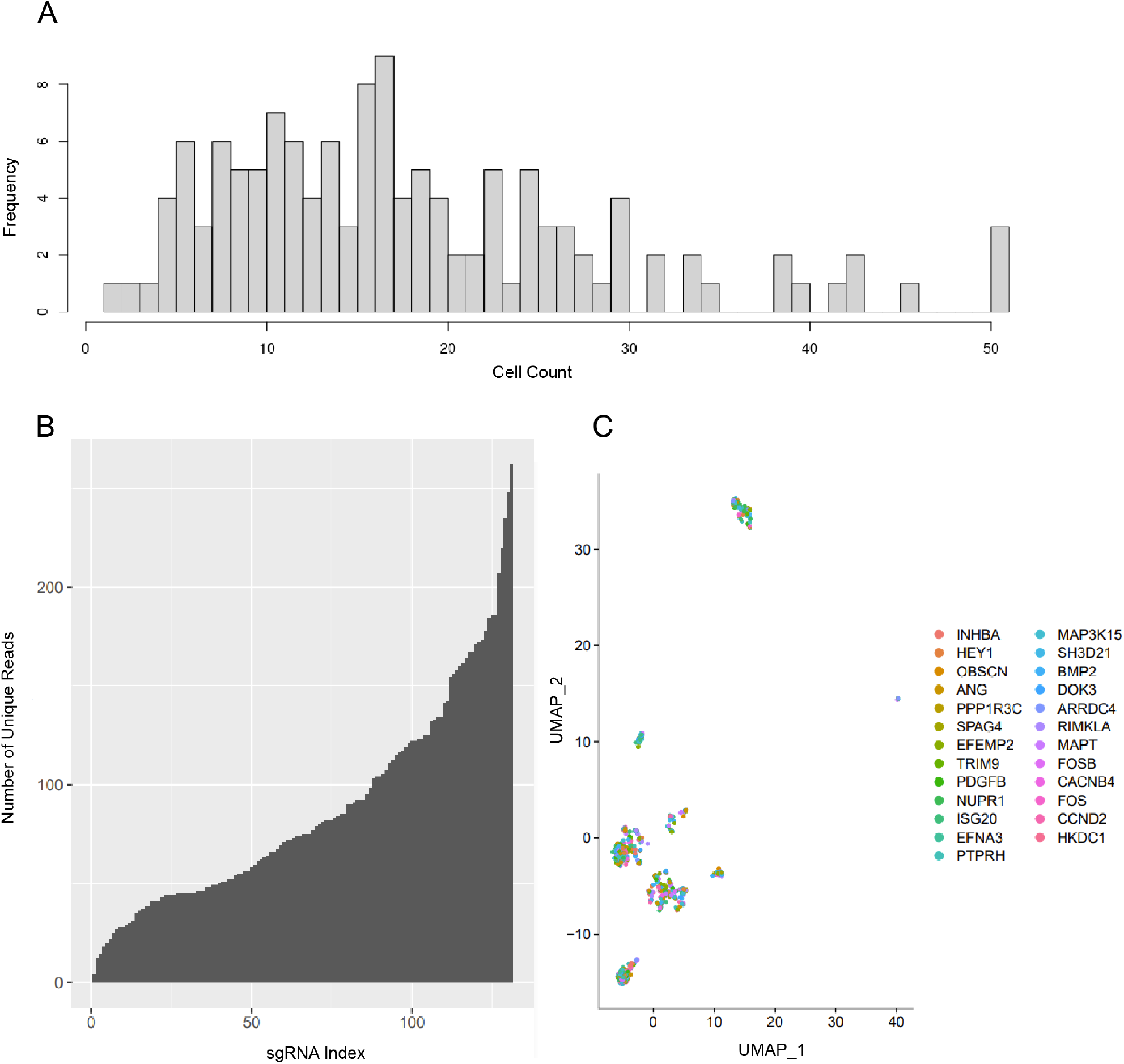
Validation of the CRISPRa experiments. (A) Cell count histogram for each sgRNA. (B) Number of reads aligned to each sgRNA. (C) A UMAP visualization of cell clusters. The colors present the sgRNA targeting indicated genes detected in the cell.

Using the in-house data, we tested the performance of our six CRISPRa prediction models. The factual and predicted fold changes of the tested genes were overall well correlated in all the models (Table 1). Particularly, we focused on the sgRNAs whose targeted genes harbored expression change within lower 25% or upper 25% in the test dataset. The predicted scores for the sgRNAs whose targets were highly induced in our experiments were significantly higher than those for the sgRNAs whose targets were barely induced (Figure 5). These data provided experimental evidence that our prediction models worked very well at least in K562 cells. Interestingly, the Bagging Regressor, Random Forest Regressor and Ada Boost Regressor models outperformed the other models (Figure 5).

**Table 1:**
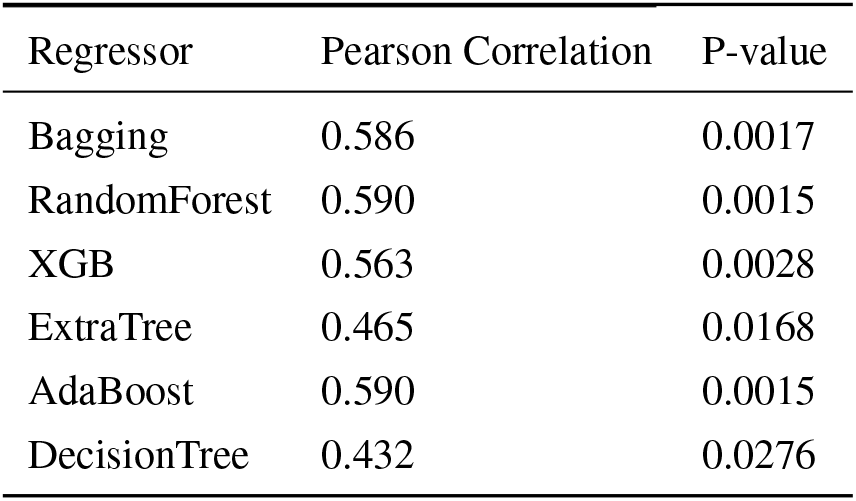
Correlation Test between the Predicted Scores and Test Dataset.

**Figure 5:**
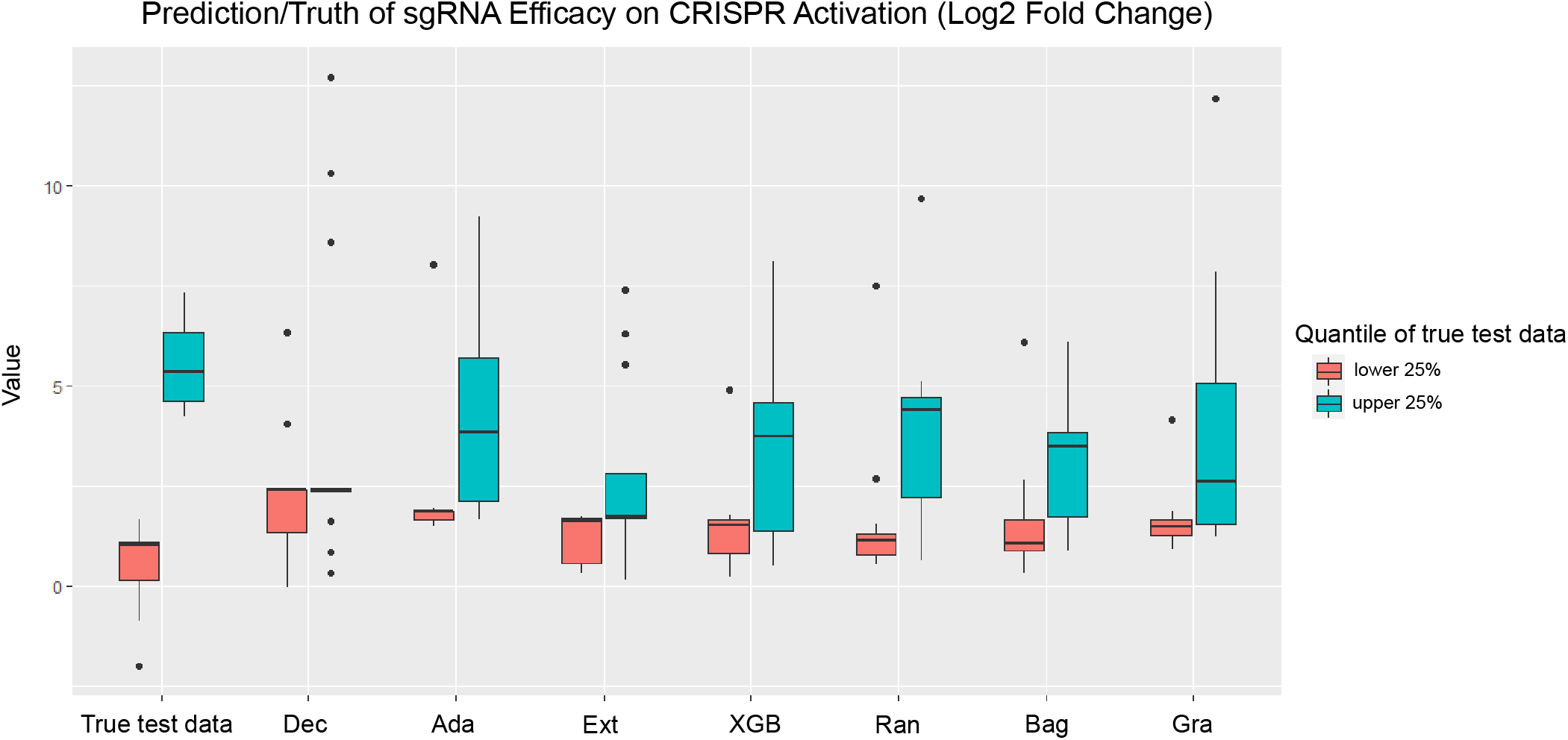
Model performance on test data. Boxplots of predicted scores of indicated sgRNAs using different models.

### 3.5 User-friendly web-tool of CRISPRa/i efficacy prediction

Finally, we designed a web-based user interface of our models and deployed it online for public access (crispr-ai.net). We named the tool “CRISPR-A-I”, which provides two function modules: 1) Activity prediction and 2) sgRNA design. In the “activity prediction” function module, users input the genomic coordinate(s) of the custom-designed sgRNA and receive the quantitative and visual epigenetic features around the region in selected cells and predicted log2 fold change of the target gene (Figure 6). In the “sgRNA design” module, our tool will design the sgRNAs for the users, which potentially perform the best in activation or interference according to users’ needs (Figure 6). Currently, four human cell lines and two mouse cell lines are available in our tool.

**Figure 6:**
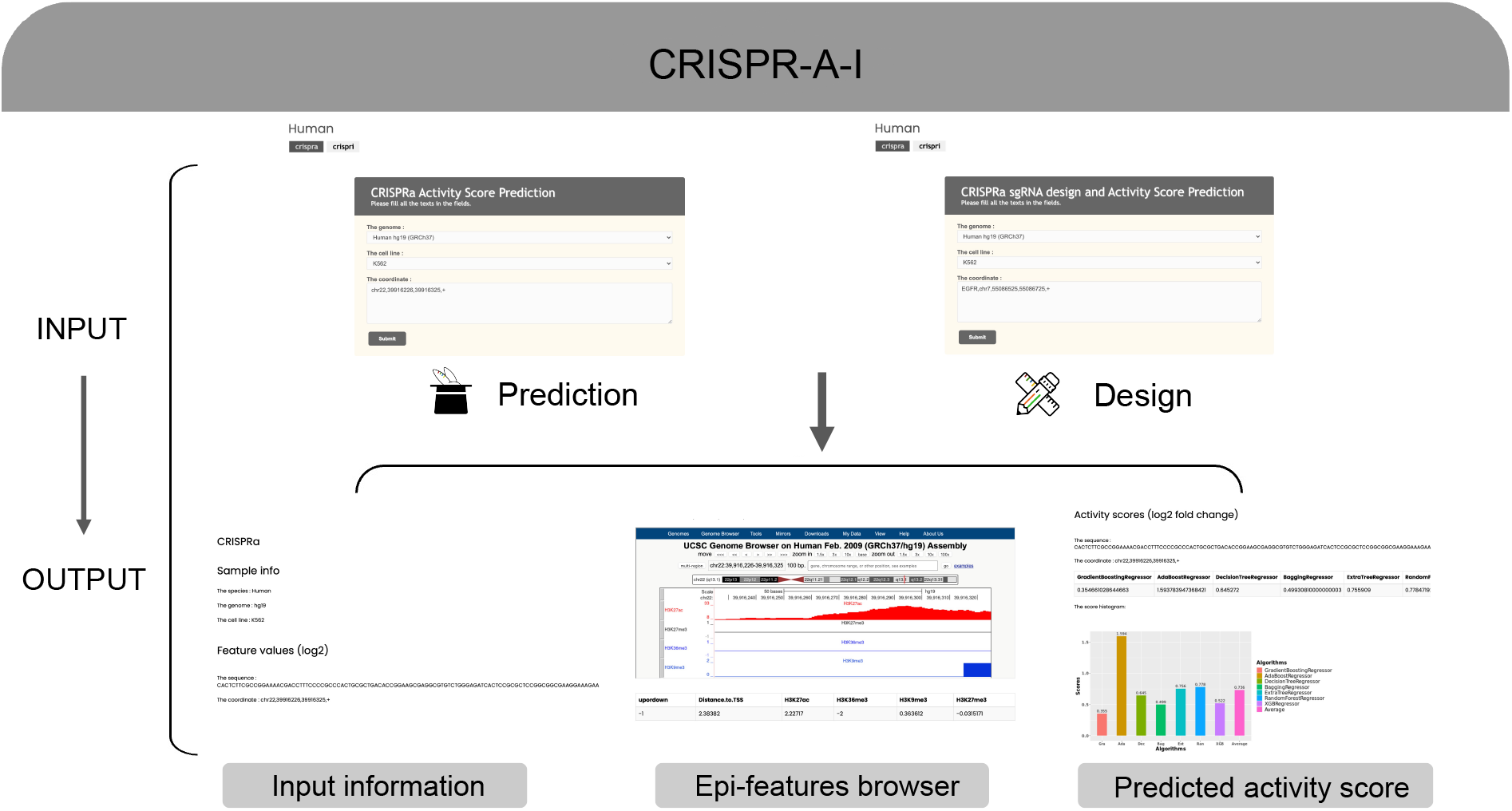
User interface of CRISPR-A-I webtool. Two modules are designed for the function of CRISPRa/i efficacy prediction of customer-defined sgRNAs and sgRNA design for customer-defined genes. The users provide genomic coordinates of the sgRNA(s) or their target gene in the two modules, respectively. The detailed description of sgRNA, visual epigenetic features in UCSC browser and the result of prediction are presented to the user eventually.

## 4 Discussion

In this study, we successfully built mathematic models as well as a web-tool predicting the efficacy of CRISPRa/I, a genetic tool extensively used by researchers in many fields. More importantly, our models were cross-validated by the experiments in our own hands. However, there are still several shortcomings in the current study. Firstly, due to the limited train dataset, it is less possible to use advanced and complex deep learning models, such as Multi-layer Perceptron Regressor (Csépe et al., 2014). A larger train dataset containing thousands of or more sgRNAs would be ideal for building a more accurate model. From the machine learning perspective, we used GC content, distance-to-TSS, and epigenetic features derived from all the available histone modification ChIP-Seq and DNase-Seq data in ENCODE as input features. However, other features, like DNA methylation may also contribute to the efficacy of CRISPRa/i. Incorporation of more features may further increase the model predicting accuracy. Moreover, the train set and our cross-validation data were both generated in K562 cells. The model in other type of cells may be subtly different and needs additional correction. Additionally, only six cell lines are selectable in our current version of tool. We will integrate more cell lines/types in the near future.

## 5 Acknowledgement

We acknowledge the ENCODE Consortium (http://www.encodeproject.org) and the laboratories generating datasets: Bradley Bernstein (Broad), John Stamatoyannopoulos (UW), Peggy Farnham (USC), Bing Ren (UCSD), and Michael Snyder (Stanford). We also thank Zhang Feng lab for providing the python code for sgRNA design.

## 6 Supplementary Information

The train data were downloaded from the ENCODE portal with the following identifiers:

ENCSR000EOT, ENCFF425WDA, ENCFF205FNC, ENCSR000APC, ENCFF840EXN, ENCFF289NOY, ENCSR668LDD, ENCFF572OAS, ENCFF407UPO, ENCSR000AKU, ENCFF804NBY, ENCFF681JQI, ENCSR000AKS, ENCFF213ZYR, ENCFF798WDB, ENCSR000AKT, ENCFF408XXB, ENCFF475HIM, ENCSR000AKV, ENCFF355NBA, ENCFF501UKW, ENCSR000AKX, ENCFF726BNA, ENCFF450IFI, ENCSR000AKP, ENCFF384ZZM, ENCFF070PWH, ENCSR000AKR, ENCFF303VNU, ENCFF190AQC, ENCSR000DWB, ENCFF481KZO, ENCFF162YFK, ENCSR000APD, ENCFF171KRI ENCFF294MOL, ENCSR000APE, ENCFF038JTR, ENCFF805FLY, ENCSR000EWB, ENCFF162VJX, ENCFF676ORH, ENCSR000EMU, ENCFF923SKV, ENCSR571IIS, ENCFF384DUK, ENCSR019SQX, ENCFF861OUG, ENCFF255JOI, ENCSR814XPE, ENCFF539XVD, ENCFF195GST, ENCSR631RJR, ENCFF568HLK, ENCFF508ASM, ENCSR000ANA, ENCFF594OXW, ENCFF111BKP, ENCSR322MEI, ENCFF385FLC, ENCFF921FHX, ENCSR952GVX, ENCFF912QYY, ENCFF330CZC, ENCSR728SZE, ENCFF157VRW, ENCFF594HBW, ENCSR000ANP, ENCFF948TVI, ENCFF661NZB, ENCSR925LJZ, ENCFF830VSZ, ENCFF554MWA, ENCSR000ANB, ENCFF004BEE, ENCFF913VVW, ENCSR301HRV, ENCFF212WAG, ENCFF147KIO, ENCSR000APZ, ENCFF616AXS, ENCFF007ZNX, ENCSR216OGD, ENCFF431KSJ, ENCFF767TKI, ENCSR000ALU, ENCFF306KIP, ENCFF228TCH, ENCSR477RTP, ENCFF869SQU, ENCSR124DYB, ENCFF668UCE, ENCFF200MQX, ENCSR087PFU, ENCFF823PVZ, ENCFF822BRL, ENCSR831JSP, ENCFF307BHB, ENCFF064SDE, ENCSR672XZZ, ENCFF398ZEA, ENCFF186BAJ, ENCSR219MYH, ENCFF051ZKE, ENCFF581JOH, ENCSR804SRO, ENCFF991DZC, ENCFF487XYM, ENCSR002YRE, ENCFF134XAW, ENCFF843EAG, ENCSR437ORF, ENCFF331WAO, ENCFF293LEM, ENCSR713QLX, ENCFF345ZBR, ENCFF532SWS, ENCSR055ZZY, ENCFF795YBZ, ENCFF793GBP, ENCFF067PJK, ENCSR431UUY, ENCFF502XME, ENCFF236RKN, ENCSR000CMW, ENCFF178GUI, ENCFF562VBE, ENCSR000CDE, ENCFF366OCH, ENCFF827TGA, ENCSR000CFO, ENCFF327OWZ, ENCFF496AXW, ENCSR000CBF, ENCFF348HPD, ENCFF121EWQ, ENCSR000CBG, ENCFF889JLT, ENCFF534ZAV, ENCSR000CGS, ENCFF092IFY, ENCFF097JSP, ENCSR000CFZ, ENCFF253YAQ, ENCFF943EUG, ENCSR000CFN, ENCFF481UTW, ENCFF513YXS, ENCSR000CEV, ENCFF807FDO, ENCFF360MLB, ENCSR000CES, ENCFF555THK, ENCFF244EVO, ENCSR000CEW, ENCFF381FVM, ENCFF592UZE, ENCSR000CEX, ENCFF977DLH, ENCFF561PGQ, ENCSR000CEU, ENCFF285VBN, ENCFF508SAM, ENCSR000ADT, ENCFF334INY, ENCFF165HOE, ENCSR000CER, ENCFF628LYK, ENCFF451SWK, ENCSR275ICP, ENCFF291ZMA, ENCFF206ALY, ENCSR265EDV, ENCFF981YBO, ENCSR043VGU, ENCFF190YKE, ENCFF955YGZ, ENCSR276HBK, ENCFF808UTB, ENCFF233ONR, ENCSR730POX, ENCFF905XXA, ENCFF431FYY, ENCSR879TAS, ENCFF790QRR, ENCFF994HSC, ENCSR729NVN, ENCFF932QYC, ENCSR876RGF, ENCFF775ONB, ENCFF930EEK, ENCSR324UNL, ENCFF882TNP, ENCFF748IXF, ENCSR188GJE, ENCFF878JJF, ENCFF494ABZ, ENCSR972SMV, ENCFF982QAB, ENCSR792GCH, ENCFF785XRM, ENCFF218GGH.

